# Brood division in a marsh-dwelling bird and its relation with the increase in offspring survival, acceleration of renest, and reduced competition for food resources

**DOI:** 10.1101/2023.10.09.561561

**Authors:** Giovanna Sandretti-Silva, L. Teixeira, C. Golec, R. Belmonte-Lopes, D. D. Sobotka, B. L. Reinert, M. A. Pizo, M. R. Bornschein

## Abstract

Parental care involves strategies that increase the offspring’s survival. In some birds, each adult only feeds one young (brood division). Six hypotheses explain the functions of brood division: 1) reduced predation; 2) selection of a young by the parents; 3) selection of an adult by the young; 4) reduction of offspring competition; 5) efficiency improvement in parent-offspring interactions; and 6) social specialization. We studied the reproductive behavior and nestling morphology of the species to evaluate the possible functions associated with the brood division in *Formicivora acutirostris*, a bird of tidal marshes of southern Brazil. Females lay two eggs, but nests with one nestling may occur. We monitored 899 nests, nestlings, and parental care of young until they reached independence. Brood division occurred when the fledglings left the nest and continued until their independence. We did not observe a difference in the frequency with which sex feed single fledglings’ clutches. The fledglings remain in separated areas (*pro* hypothesis #1). There is a sex bias in the division (*pro* hypothesis #2). Adults fed the siblings equally (*contra* hypothesis #3). There is no aggression between siblings (*contra* hypothesis #4). Each fledgling remained close to their adult and responds only to it (*pro* hypotheses #5 and #6). We propose that brood division in the species may 7) reduce drowning of fledglings, 8) spatially divide food consumption, and 9) allow concomitant reproduction. We emphasize flooding as a structuring force for *F. acutirostris*, which raises concerns about the effects of climate change on this species.

## Introduction

For most birds, parental care after birth is carried out by both sexes (Trémont and Ford 2000; Cockburn 2006). In some species, each adult assumes exclusive care of only one young (e.g., Wilson and Kikkawa 1988; Draganoiu et al. 2006; Vega et al. 2007; Tarwater and Brawn 2008; Watson et al. 2012). This division usually happens after the fledgling leaves the nest, but it can also occur in the nestling phase (Tarwater and Brawn 2008). Six hypotheses were proposed to explain the brood division and its associated functions: (1) reduction in brood predation by segregating fledglings spatially in the context of a high chance of mortality, (2) selection of a specific young based on sex or size, (3) selection of an adult with better skills in food provisioning, (4) reduction in the conflict between siblings to homogenize their ability to get food and reduce aggression between them, (5) improvement of the effectiveness of parental care on a specific fledgling, and (6) improvement in the efficiency of the interaction between adults and fledglings, which is called social specialization (Tarwater and Brawn [2008] and a review of the literature included).

A family of birds in which brood division occurs is Thamnophilidae, a Neotropical family consisting of 234 species of mostly forest insectivorous birds (Winkler et al. 2020). Almost all Thamnophilidae are monogamous and form permanent pairs that defend their territories together throughout the year (Zimmer and Isler 2003; see Willis 1978; Skutch 1996). Both male and females are responsible for the care of nestlings and fledglings (e.g., Skutch 1946; Willis and Oniki 1972; Greenberg and Gradwohl 1983), but each adult feed only one young, ignoring the other or feeding it only sporadically (e.g., Skutch 1969; Oniki 1975; Tarwater and Brawn 2008). In *Formicivora erythronotos* the division was defined by the adult who witnessed a particular fledgling leaving the nest (Mendonça 2001), while in *Thamnophilus atrinucha* the male selected the heaviest nestling or the first fledgling to leave the nest (Tarwater and Brawn 2008). In *T. atrinucha*, brood division is mandatory to avoid predation (hypothesis #1) and by social specialization (hypothesis #6), even in broods of only one young (Tarwater and Brawn 2008). This is the only species of Thamnophilidae tested for regularity and associated functions of brood division.

Studies of parent behavior in contrasting environmental contexts provide an opportunity to explore how different habitats can shape parental care strategies (Tarwater and Brawn 2008) in response to variables that affect both offspring and parent survival. In this way, only two species of Thamnophilidae live exclusively in marshes (Bornschein et al. 2015; See also Zimmer and Isler 2003: 492), the Parana antwren *Formicivora acutirostris* (Bornschein et al. 1995) and Sao Paulo antwren *F. paludicola* (Buzzetti et al. 2013). The first species occurs in coastal marshes flooded daily by tides (see habitat details in Reinert et al. [2007]), which can put a toll on reproduction due to the frequent drowning of nests, with consequences for parent care (Reinert et al. 2007, 2012). In the present study, we investigated brood division and explored the possible hypotheses related to its associated functions to increase productivity in *F. acutirostris*, a marsh-dwelling bird endemic to the Brazilian Atlantic Forest (Brooks et al. 1999).

## Methods

### Study species

*Formicivora acutirostris* is socially monogamous, with both sexes forming long-lived pairs that participate in breeding and territory defense activities throughout the year (Reinert et al. 2012; Bornschein et al. 2015). The pair builds a half-cup nest where two eggs are laid and a maximum of four young can be produced per breeding season, which extends from August to February (Reinert et al. 2012; Bornschein et al. 2015). Nests can be lost to flooding, predation, and tipping, leading to up to eight nesting attempts per pair in a breeding season (on average, four per pair; Reinert 2008; Reinert et al. 2012).

### Study areas

The study region is the Ramsar Site Guaratuba at Guaratuba Bay, municipality of Guaratuba, southern Brazil, where we worked in Jundiaquara Island (2006 – 2022; *c.* 25.873611°S, 48.758889°W; 11.50 ha), Claro River (2007 – 2022; *c.* 25.874444°S, 48.762222°W; 8.73 ha), Folharada Island (2009 – 2022; *c.* 25.866111°S, 48.723056°W; 16.30 ha) and Lagoa do Parado (2012 – 2013; *c.* 25.743333°s, 48.714722°W; 9.87 ha). From January 2006 to May 2008 fieldwork was conducted every day during the breeding season, and for 6 – 8 days a month in non-breeding seasons. From June 2008 to March 2022 we worked 3 – 8 days per month in the field. See Reinert et al. (2007) for more details on the study areas.

### Field procedures

We explored the brood division and its associated functions, based on the hypotheses from Tarwater and Brawn (2008). To investigate brood division, we recorded nestling and fledgling care, adult behavior towards fledglings, and parental care in single-fledgling broods. For the associated functions, we observed egg-laying interval and incubation, size, plumage, and sex of nestling, nestling behavior, nestling care, departure from the nest, fledgling care, adult behavior towards fledglings, fledgling survival, and brood division period (Table 1).

**Table 1.** Support procedure used to assess the applicability of the hypothesis that explain the brood division and its associated functions in the Paraná Antwren (*Formicivora acutirostris*).

| Hypothesis | Observed traits | Accepted | Rejected |
| --- | --- | --- | --- |
| 1. Predation reduction | Fledgling care and survival; adult behavior towards fledglings | Spatially divided brood; present defensive behaviors of parents; predation risk; low fledgling mortality | Non-spatially divided brood; absent defensive behaviors of parents; low predation risk; high fledgling mortality |
| 2. Selection of a young | Egg laying interval; frequency of incubation, even during the egg laying period, to assess a prior differentiated level of parental care; size, plumage, and sex of nestlings | Distinct fledglings; selection of fledglings based on their size and/or sex | Identical fledglings; random selection of fledglings |
| 3. Selection of an adult | Frequency of incubation, even during the egg laying period, to assess to assess a prior differentiated level of parental care; nestling care; departure from the nest | Differential nestling care between parents; separated nest departure following an adult | Identical nestling care between parents; joint nest departure |
| 4. Offspring<br>conflict reduction | Nestling care and behavior | Brood division during nestling phase,<br>or aggression between nestlings before<br>brood division | Brood division after nestling phase,<br>and no aggression between nestlings<br>before brood division |
| 5. Parental care<br>improvement | Fledgling care | Fledgling closest to its adult;<br>improved food efficiency | Fledgling random positioned; food<br>efficiency not improved |
| 6. Interaction<br>improvement | Fledgling care, adult behavior towards<br>nestlings, brood Division period | Long-term brood division; improved<br>recognition between adult and<br>fledgling | Short term brood division; improved<br>recognition between adult and<br>fledgling |

<sup>a</sup> According to Tarwater and Brawn [2008] and a review of the literature included

We marked all territorial individuals of the study sites with unique combinations of colored bands. To assist in the monitoring of breeding pairs and record parental care, we determined their territories in all areas sampled by recording the routes of each member of a pair within a grid marked with numbered stakes fixed every 25 m, or with GPS. See Bornschein (2013) for further details on the determination of territories.

We monitored monthly all the territories in each study area by following marked individuals for the observation of reproductive activities. We georeferenced all the nests found and monitored their fates returning every other day to avoid trampling the fragile vegetation which might call the attention of predators. To detect reproductive activities in the egg and nestling phase, we made direct observations at 4 m from the nests (50 sessions totaling 56 h 19 min), in tents at 2 m from the nests (252 sessions, 323 h 23 min), and by installing camcorders positioned at 1.5 m from the nests (131 sessions, 130 h 51 min), alternating the sessions throughout the day. We defined the incubation period as the time between the laying of the second egg and the birth of the last hatching (modified from Nice [1954]). Full incubation shifts represent the incubation time from the arrival of an adult to the nest until its exit. Incomplete incubation shifts represent the incubation time considering the observation of the individual who was already incubating when our focal observations began.

Nestlings were only manipulated for banding and biometrics when they were about to leave the nest. We captured by hand fledglings that had recently left the nest, which were weighed (10 g dynamometer, 0.1 g scale) and measured (digital plastic caliper, 0.1 mm scale). We monitored fledglings until they became independent, which occurs between 7 – 9 weeks of life (Reinert 2008), recording which adult took care of them, and observed interactions such as feeding, calling, and moving together. We confirmed the sex of some young based on the plumage succession by monitoring 12 males and 11 females until they reach adult plumage, with an accentuated sexual dimorphism. It was not possible to record data blind because our study involved focal animals in the field.

### Data analysis

We performed a Shapiro-Wilk test to verify the normality of the data on food provisioning to nestlings before performing a paired t-test to investigate for differences in the frequency of food provisioning between parents. We assessed whether each parent cared for male or female young at random (expected ratio of 1:1) by performing a chi-square test with α = 5%. The same test was used to evaluate if male or female parents preferentially care for the young in single-fledgling broods, to determine if he was cared by the male or female randomly (expected ratio of 1:1), also performing a chi-square test (α = 5%). Due to differences in data precision, we performed this test considering the cases in which we confirmed the reduction of the brood in the nest and considering the cases in which we detected only one fledgling cared by the adult, without knowing if there was a reduction of the brood in the nest or if two fledglings left the nest and one died unnoticed. To compare the proportions of confirmed and unconfirmed cases, we performed a Fisher’s exact test (α = 5%).

## Results

We monitored 12 – 41 territories by year, totaling 490 territories monitored for 12 months (Table S1). A total of 899 nests were found, of which 738 were concluded. We monitored 271 juveniles that achieved independence, of which 24 marked as nestlings were recruited in monitored territories (Table S2).

### Evaluating the selection of a young by the parents (hypothesis #2)

#### Egg laying interval, incubation, and the size of nestlings

The interval of days between the laying of the first and second eggs was recorded in 29 nests: in 12, the eggs were laid without interval; in 14, there was a daybreak; and in three two days break. Laying intervals were still estimated at 16 nests: in six, the posture would have been done in consecutive days; in nine, there would have been a daybreak; and in a single nest, there would have been two days break. The average day interval between the laying of the two eggs, considering recorded and estimated data, was 1.69 day (± 0.63 day).

The incubation of the first egg is irregular. Out of 17 monitored nests, in nine both adults incubated the first egg, in four only the males incubated, and in the other four nests, only the females incubated. Additionally, the incubation time of the first egg is short, on average 12.5 min (± 8.70 min; n = 15 nests) considering the incomplete shifts, and 12.3 min (± 9.00 min; n = 8 nests) for complete shifts. From the laying of the second egg, the incubation shifts became longer, lasting on average 32.7 min when females incubate, and 46.1 min when males incubate (Table 2). The incubation period last 14 – 18 days (*x* = 16.0 days ± 1.07 day; n = 18 nests). Probably because the incubation of the first egg is irregular, the nestlings of *F. acutirostris* have similar sizes (Table 3). Eight nestlings, on the day they left the nests, had similar body mass, ranging from 6.8 – 7.5 g (*x* = 7.3 g ± 0.36 g).

**Table 2.**
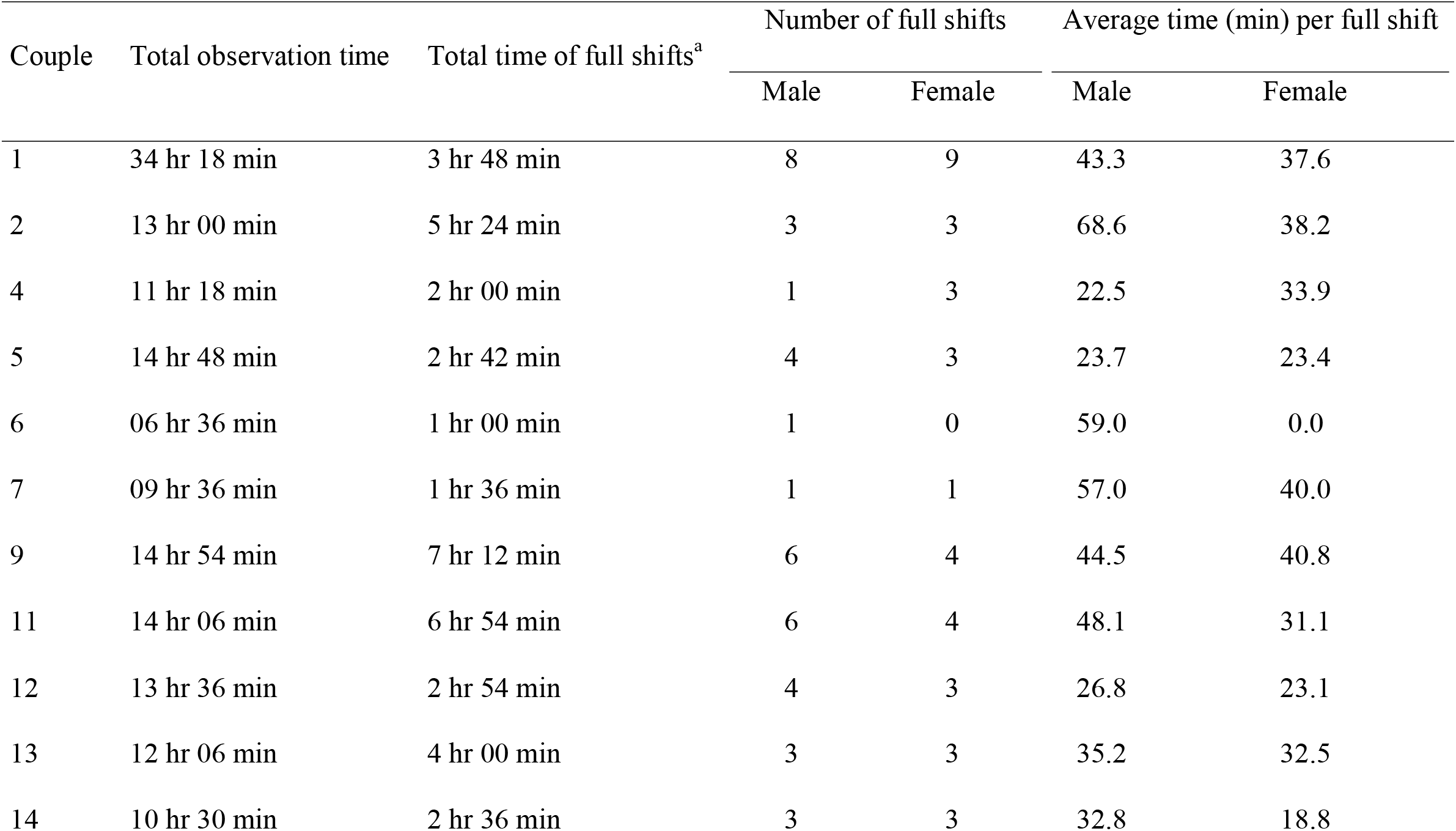
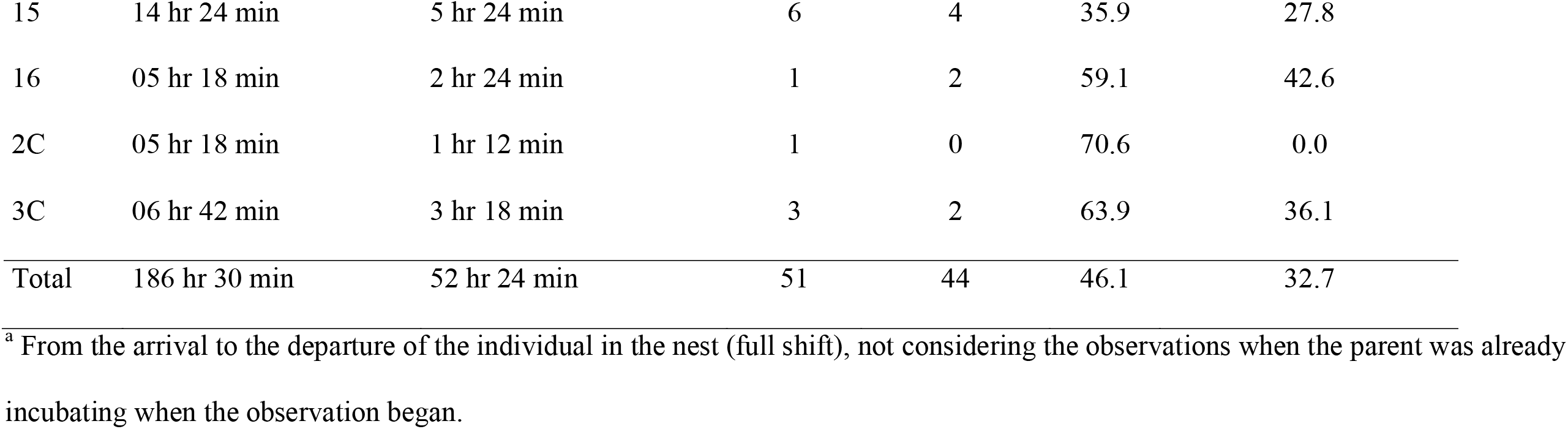
Duration of full incubation shifts of *Formicivora acutirostris* in the reproductive seasons of 2006 / 2007 and 2007 / 2008 on the Jundiaquara Island and Claro River, Guaratuba Bay, Paraná, southern Brazil.

| Couple | Total observation time | Total time of full shifts<sup>a</sup> | Number of full shifts |  | Average time (min) per full shift |  |
| --- | --- | --- | --- | --- | --- | --- |
|  |  |  | Male | Female | Male | Female |
| 1 | 34 hr 18 min | 3 hr 48 min | 8 | 9 | 43.3 | 37.6 |
| 2 | 13 hr 00 min | 5 hr 24 min | 3 | 3 | 68.6 | 38.2 |
| 4 | 11 hr 18 min | 2 hr 00 min | 1 | 3 | 22.5 | 33.9 |
| 5 | 14 hr 48 min | 2 hr 42 min | 4 | 3 | 23.7 | 23.4 |
| 6 | 06 hr 36 min | 1 hr 00 min | 1 | 0 | 59.0 | 0.0 |
| 7 | 09 hr 36 min | 1 hr 36 min | 1 | 1 | 57.0 | 40.0 |
| 9 | 14 hr 54 min | 7 hr 12 min | 6 | 4 | 44.5 | 40.8 |
| 11 | 14 hr 06 min | 6 hr 54 min | 6 | 4 | 48.1 | 31.1 |
| 12 | 13 hr 36 min | 2 hr 54 min | 4 | 3 | 26.8 | 23.1 |
| 13 | 12 hr 06 min | 4 hr 00 min | 3 | 3 | 35.2 | 32.5 |
| 14 | 10 hr 30 min | 2 hr 36 min | 3 | 3 | 32.8 | 18.8 |
|  |  |  | Male | Female | Male | Female |
| 15 | 14 hr 24 min | 5 hr 24 min | 6 | 4 | 35.9 | 27.8 |
| 16 | 05 hr 18 min | 2 hr 24 min | 1 | 2 | 59.1 | 42.6 |
| 2C | 05 hr 18 min | 1 hr 12 min | 1 | 0 | 70.6 | 0.0 |
| 3C | 06 hr 42 min | 3 hr 18 min | 3 | 2 | 63.9 | 36.1 |
| Total | 186 hr 30 min | 52 hr 24 min | 51 | 44 | 46.1 | 32.7 |

<sup>a</sup> From the arrival to the departure of the individual in the nest (full shift), not considering the observations when the parent was already incubating when the observation began.

**Table 3.** Measurements (mm) and body mass (g) of nestlings (indeterminate sex) of *Formicivora acutirostris* with 7 – 11 days of life (X = 8.5 days) in the reproductive seasons of 2006 / 2007 and 2007 / 2008 on the Jundiaquara Island, Guaratuba Bay, Paraná, southern Brazil.

| Measurement | n | Range | Mean and standard deviation |
| --- | --- | --- | --- |
| Body mass | 16 | 6.7 – 8.6 | $7.5 \pm 0.52$ |
| Culmen | 13 | 7.2 – 10.0 | $8.7 \pm 0.90$ |
| Distal margin from nostril to tip of beak | 16 | 3.0 – 6.5 | $5.0 \pm 0.83$ |
| Wing | 16 | 32.6 – 39.1 | $35.6 \pm 2.03$ |
| Tarsus | 16 | 21.2 – 25.4 | $23.8 \pm 1.22$ |

#### Sex-related plumage and care

Nestlings had a similar appearance, with only subtle sexual dimorphism made by the extension and coloring of the apex of the alula and primary coverts. These sex-related features emerge from pins after some days, but always before young departure from nests. In males, the apex has white spots, while in females it is endowed with whitish spots, ranging from almost imperceptible whitish edges to large spots as that of males (Figure 1).

**Figure 1.**
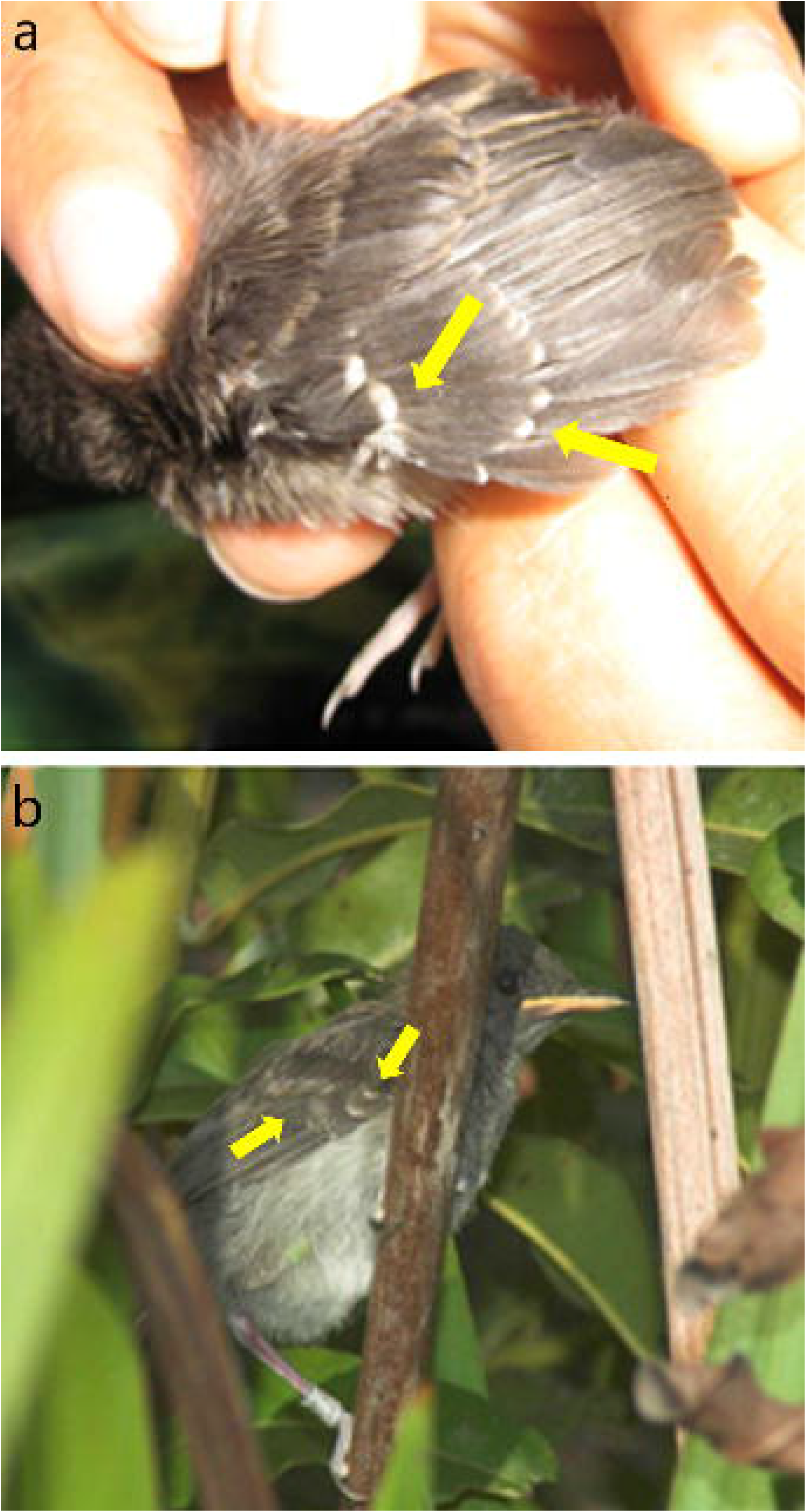
Sexual dimorphism in fledglings of *Formicivora acutirostris*, Jundiaquara Island, Guaratuba Bay, Paraná, southern Brazil. (a) male (Fi5) nine days old when he left the nest. (b) female (Fi15) with 22 days of life. Note the white apices of the alula and upper primary of the male against the more limited and whitish apices of the female. We confirmed the sex determination by seeing these individuals months later. Photos: Ricardo Belmonte-Lopes.

The young replaced all the upper and lower contour feathers and upper and lower coverts of the tail and wings, with exception of primary coverts and alula (the feathers that present sexual dimorphism since nestlings), before reaching independence. They had then a female plumage, presenting, again, subtle sexual dimorphism, with males having the breast less striated white and black, typical of the females, and rather with streaks or gray spots amid white streaks. The young retain the primary coverts and alula, developed when they were in the nest, until they reach independence.

We were able to confirm the sex of 23 marked fledglings by observing them at least when they were eight months old. The following correspondence occurred between the sex of the fledgling and the respective adult who cared it: adult males caring for male fledglings (n = 11), adult males caring for female fledglings (n = 5), adult females caring for female fledglings (n = 6), and adult females caring for male fledgling (n = 1). Of these six cases of absence of correspondence between the sex of the adult and the respective fledgling, in two it would not be possible to obtain it because the brood contained siblings of the same sex (2 males in one nest and 2 females in another). Unfortunately, we were unable to confirm the sex of the second fledgling of the brood in the remaining four cases of no correspondence. There is a correspondence between the sex of the adult and its fledgling (X² = 5.26; *p* = 0.02; df = 1).

### Evaluating the selection of an adult by the young (hypothesis #3), and the reduction of offspring competition (hypothesis #4)

#### Nestling care, nestling behavior, and departure from nests

The frequency of food provisioning to nestlings was similar between the adults (t = 0.992; df = 12; *p* = 0.34; Table 4). We did not observe aggression between nestlings in observations near nests or in filmed nests. Nestlings leave the nests together when they were 7 – 11 days old (X = 9 days).

**Table 4.** Age (days) of fledglings of *Formicivora acutirostris* at the day before they were forced to abandon the territory by the parents in the reproductive seasons of 2006 / 2007 and 2007 / 2008 on the Jundiaquara Island, Guaratuba Bay, Paraná, southern Brazil.

| Fledgling | Couple | Brood<sup>a</sup> | Parent sex | Observation days | Age |
| --- | --- | --- | --- | --- | --- |
| Fi6 | 11 | 1<sup>st</sup> | male | 4 | 59 |
| Fi7 | 11 | 1<sup>st</sup> | female | 6 | 58 |
| Fi8 | 15 | 1<sup>st</sup> | male | 8 | 54 |
| Fi9 | 7 | 1<sup>st</sup> | female | 5 | 49 |
| Fi10 | 7 | 1<sup>st</sup> | female | 6 | 71 |
| Fi11 | 5 | 1<sup>st</sup> | male | 7 | 48 |
| Fi15 | 10 | 1<sup>st</sup> | female | 12 | 62 |
| Fi17 | 6 | 2<sup>nd</sup> | male | 11 | 61 |
| Fi19 | 11 | 2<sup>nd</sup> | male | 14 | 49 |

<sup>a</sup> The indication of a second brood means that the pair has produced fledglings in a previous attempt in the same breeding season.

### Evaluating the reduction of predation (hypothesis #1), improvement of the efficiency of parent-offspring interactions (hypothesis #5), and social specialization (hypothesis #6)

#### Duration of parental care and survival

Fledglings reach independence between 49 – 71 days of life (Table 5). Of 214 fledglings that we monitored since they left the nests, 153 (71.5%) reached independence. Of the remainders, 17 died on the day they left the nest, 10 died after that, and 34 died on an unknown day (Figure 2). We detected the cause of death of nine fledging on the day they left the nests: three were preyed and seven drowned. Four of them could not stand on the vertical vegetation in which they were and slipped falling into the water, and three fell into the water while moving amidst the vegetation. Drowning not only involved the fall of the fledgling in the water, but also instances in which the water reached the fledglings by the rise of the tide. Once, a fledgling flew into the mud after two ants (*Camponotus* sp.) bite. It fluttered its wings as it received food from the male, but when the tidal water reached its fingers, the fluttering of its wings splashed water on it (Figure 3A). The water level did not pass from the fingers of the fledgling, and it dried up.

**Figure 2.**
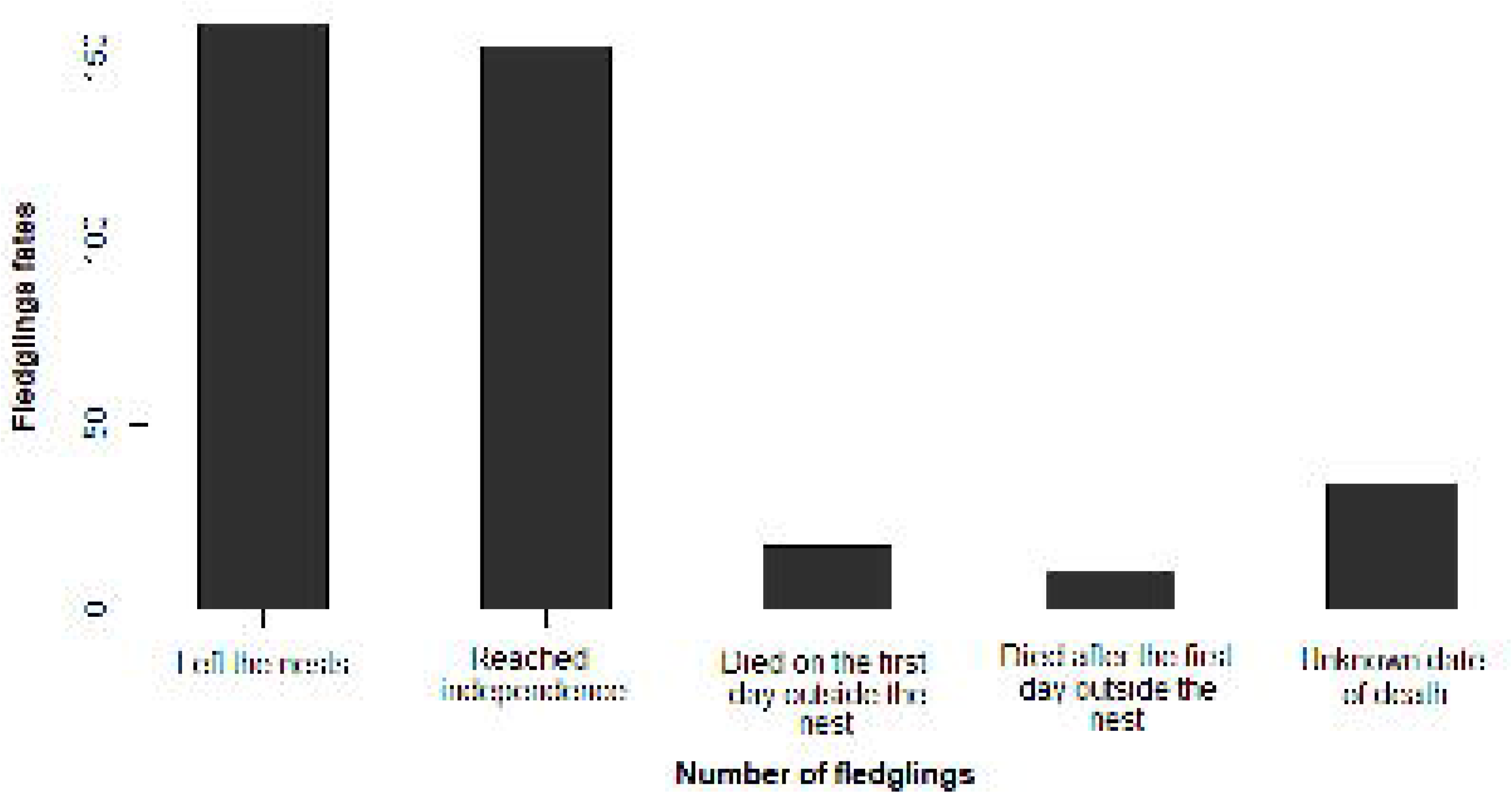
Fate of fledglings of *Formicivora acutirostris*, Guaratuba Bay, Paraná, southern Brazil

**Figure 3.**
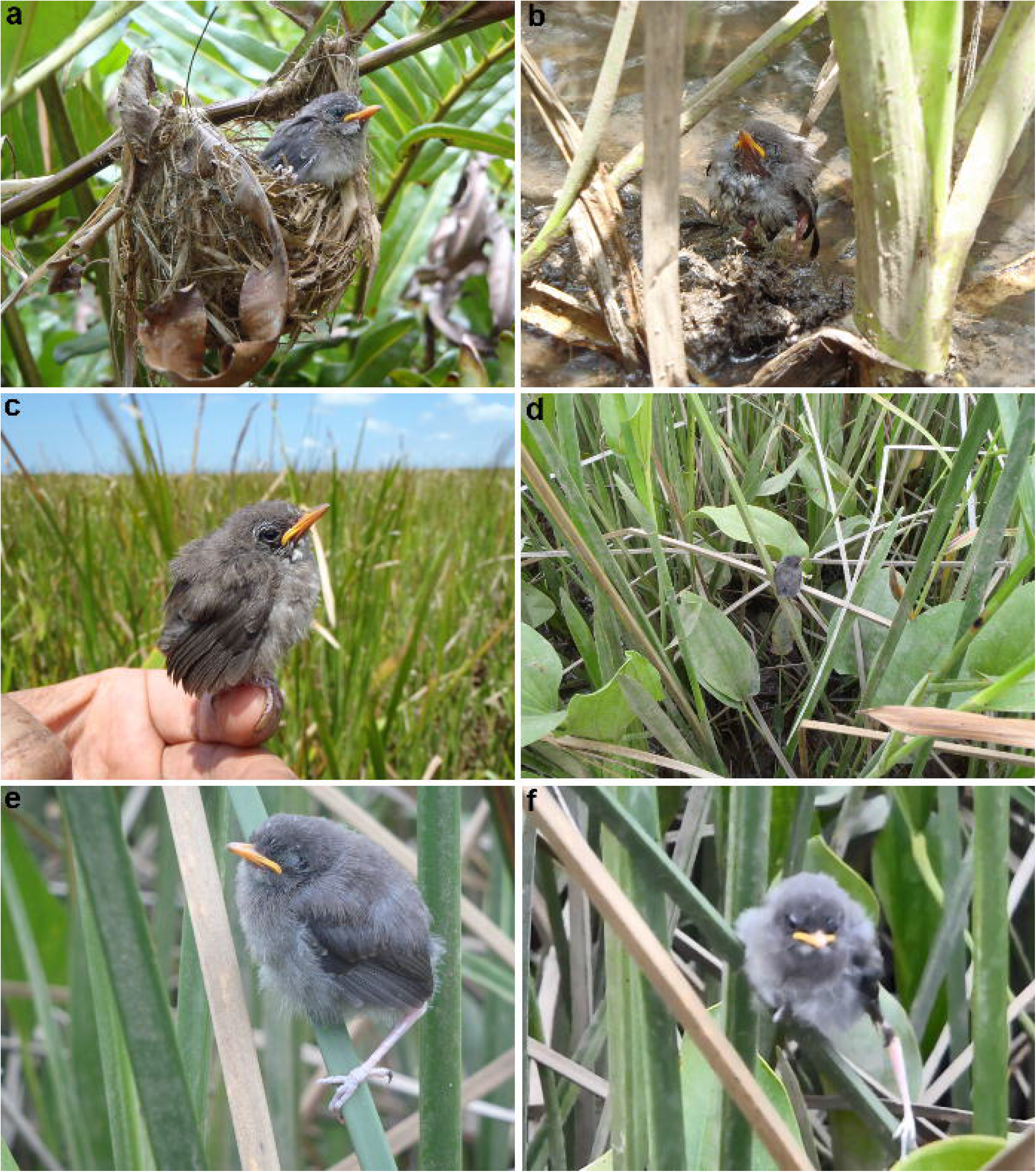
Three fledglings of *Formicivora acutirostris* at the day they left the nests, Guaratuba Bay, Paraná, southern Brazil. In (a) there is a nestling in the edge of the nest a few moments before leaving it. In (b) and (c) there is a second fledgling that flew into the mud after being stung by ants on the fingers (b) and was captured with free hands to be marked (c) after we verified that it survived the tide, which soon retreated. Notice the wet feathers by vibrating the wings as the male approached to feed it. In (d)–(f) there is a third fledgling, moving in the vegetation less than 1 m high over the substrate (d), sleeping (e), and slipping the left leg (f). Photos and snapshot of a video (f): Marcos R. Bornschein

**Table 5.** Visits from parents of *Formicivora acutirostris* to the nests during the feeding period of nestling in the reproductive seasons of 2006 / 2007 and 2007 / 2008 on the Jundiaquara Island and Claro River, Guaratuba Bay, Paraná, southern Brazil.

| Couple | Total | Total time of | Total | Total number of |  | Total number of visits with |  | Time interval (min) between |  |
| --- | --- | --- | --- | --- | --- | --- | --- | --- | --- |
|  | observation | the nest being | unattended | visits |  | food |  | visits |  |
|  | time | attended | nest time | Male | Female | Male | Female | Male | Female |
| 1 | 02 hr 12 min | 00 hr 42 min | 01 hr 30 min | 6 | 71 | 4 | 70 | 38.7 | 8.3 |
| 2 | 16 hr 54 min | 08 hr 36 min | 08 hr 18 min | 118 | 92 | 112 | 87 | 13.1 | 20.4 |
| 4 | 04 hr 06 min | 03 hr 30 min | 00 hr 36 min | 9 | 8 | 7 | 6 | 27.6 | 31.0 |
| 6 | 13 hr 30 min | 07 hr 12 min | 06 hr 18 min | 77 | 135 | 69 | 116 | 28.8 | 36.7 |
| 7 | 05 hr 00 min | 02 hr 12 min | 02 hr 48 min | 50 | 34 | 49 | 28 | 6.0 | 8.9 |
| 9 | 01 hr 18 min | 00 hr 30 min | 00 hr 48 min | 20 | 44 | 20 | 41 | 3.8 | 1.7 |
| 10 | 07 hr 12 min | 03 hr 18 min | 03 hr 54 min | 44 | 47 | 41 | 44 | 9.9 | 9.2 |
| 11 | 06 hr 06 min | 04 hr 00 min | 02 hr 06 min | 33 | 21 | 31 | 16 | 14.9 | 17.5 |
| 12 | 08 hr 30 min | 04 hr 36 min | 03 hr 54 min | 36 | 58 | 29 | 54 | 14.1 | 8.7 |
| 13 | 01 hr 18 min | 01 hr 06 min | 00 hr 12 min | 2 | 2 | 2 | 2 | 37.5 | 37.5 |
|  | observation | the nest being | unattended | visits |  | food |  | visits |  |
|  | time | attended | nest time | Male | Female | Male | Female | Male | Female |
| 15 | 09 hr 06 min | 01 hr 48 min | 07 hr 18 min | 118 | 120 | 117 | 118 | 4.7 | 4.6 |
| 16 | 05 hr 12 min | 04 hr 12 min | 01 hr 00 min | 19 | 14 | 13 | 8 | 16.3 | 22.1 |
| 2C | 07 hr 12 min | 06 hr 06 min | 01 hr 06 min | 25 | 22 | 19 | 17 | 17.2 | 19.5 |
| Total | 87 hr 36 min | 47 hr 48 min | 39 hr 48 min | 557 | 668 | 513 | 607 | 17.9 | 17.4 |

#### Adult behavior toward fledglings

From the first day outside the nests, each adult and their respective fledglings went to different areas of the territory and usually do not overlap foraging sites. Fledgling recognizes the call of its caring adult and from the second day outside the nest respond with hoots and / or moving towards him and ignoring the calls of the other adult. All fledglings that slipped close to the water level had their respective adult close to them vocalized a lot in a higher position and only silenced when the fledgling reached greater height. The time that the fledgling was left alone increases when they reach around 30 days of age and can last for up to 68 min when they are close to independence.

### Brood division

We observed brood division always on the first day after the fledglings left the nest. Each parent attended only one fledgling permanently until it reaches independence. Brood division happened even when only one nestling is produced. In this case, one adult fed the fledgling while the other did not carry out any reproductive activity. Sporadically females were observed feeding the fledgling cared by the male, but we never recorded the opposite (Table 6). However, sometimes males feed the females, which gives this “extra” food to the fledglings they care. An exception to brood division occurred in a single case when the male of a pair who had two fledglings disappeared; the female then adopted the fledgling that was under the care of the male.

**Table 6.** Number of times we have seen fledglings of *Formicivora acutirostris* being fed in the reproductive seasons of 2006 / 2007 and 2007/2008 on the Jundiaquara Island, Guaratuba Bay, Paraná, southern Brazil.

| Fledgling | Couple | Brood<sup>a</sup> | Number of 20-min<br>observation events | Number of times it was fed by |  | Average time (min) of the interval |  |
| --- | --- | --- | --- | --- | --- | --- | --- |
|  |  |  |  | parental |  | between feeds |  |
|  |  |  |  | Male | Female | Male | Female |
| Fi8 | 15 | 1<sup>st</sup> | 7 | 60 | 7 | 5.5 | 16.4 |
| Fi10 | 7 | 1<sup>st</sup> | 1 | 0 | 12 | --- | 1.7 |
| Fi11 | 5 | 1<sup>st</sup> | 3 | 62 | 6 | 1.1 | 12.2 |
| Fi15 | 10 | 1<sup>st</sup> | 7 | 0 | 17 | ? | 8.4 |
| Fi17 | 6 | 2<sup>nd</sup> | 1 | 13 | 0 | 1.5 | --- |
| Fi19 | 11 | 2<sup>nd</sup> | 8 | 88 | 1 | 2.0 | 20.0 |
| Fi20 | 11 | 2<sup>nd</sup> | 2 | 0 | 11 | --- | 4.0 |
| Fi21 | 15 | 2<sup>nd</sup> | 3 | 0 | 18 | --- | 3.3 |

<sup>a</sup> The indication of a second brood means that the couple has produced fledglings in a previous attempt in the same breeding season

### Adults that care single-fledgling broods

We monitored 13 cases in which there was brood reduction and the nest had only one nestling: in 10 the fledgling was fed by the male and in the remaining three by the female. The causes of brood reduction were: an egg fell down from inclined nests, an egg carried away by high tide, an egg thrown out of the nest by the wave of a boat, an egg rotted in the nest, and a nestling preyed upon in the nest. Of 91 single-fledgling broods, without confirmation of brood reduction, 47 were fed by the male and 44 by the female, with no difference in the propensity of any sex to care for the single fledgling (X^2^ = 0.99; *p* = 0.75; df = 1). There was no difference between the proportions of care for each sex between confirmed and unconfirmed data (*p*=0.98), therefore, in none of the conditions one sex of adult cared more in single-fledgling broods.

### Renest in cases of single-fledgling broods

In a pair in which only one member was caring for a fledgling, vocalizations that precede the copulation were constantly heard from the member that was “free”. We often observed males without fledglings starting the construction of nests and with pre-copulation behavior towards females who were caring for fledglings and who, apparently, remained indifferent to them. Without the contribution of females, these nests were not built beyond the initial stage. However, we witnessed five cases of concomitant reproductions. In four of them, the males were with a fledgling and the respective females were not, while in one case the female was with a fledgling and the male was not. These situations occurred when adults cared for single 30 – 40 days old fledglings and simultaneously had nests with eggs.

Two of the studied concomitant reproductions resulted in the only case of three broods with fledglings reaching independence observed in *F. acutirostris* (Figure 4C). On that occasion, the first concomitant reproduction happened when the male was with a fledgling, in October, and the second when the female was with a fledgling, in December. The third fledgling was cared by the male.

**Figure 4.**
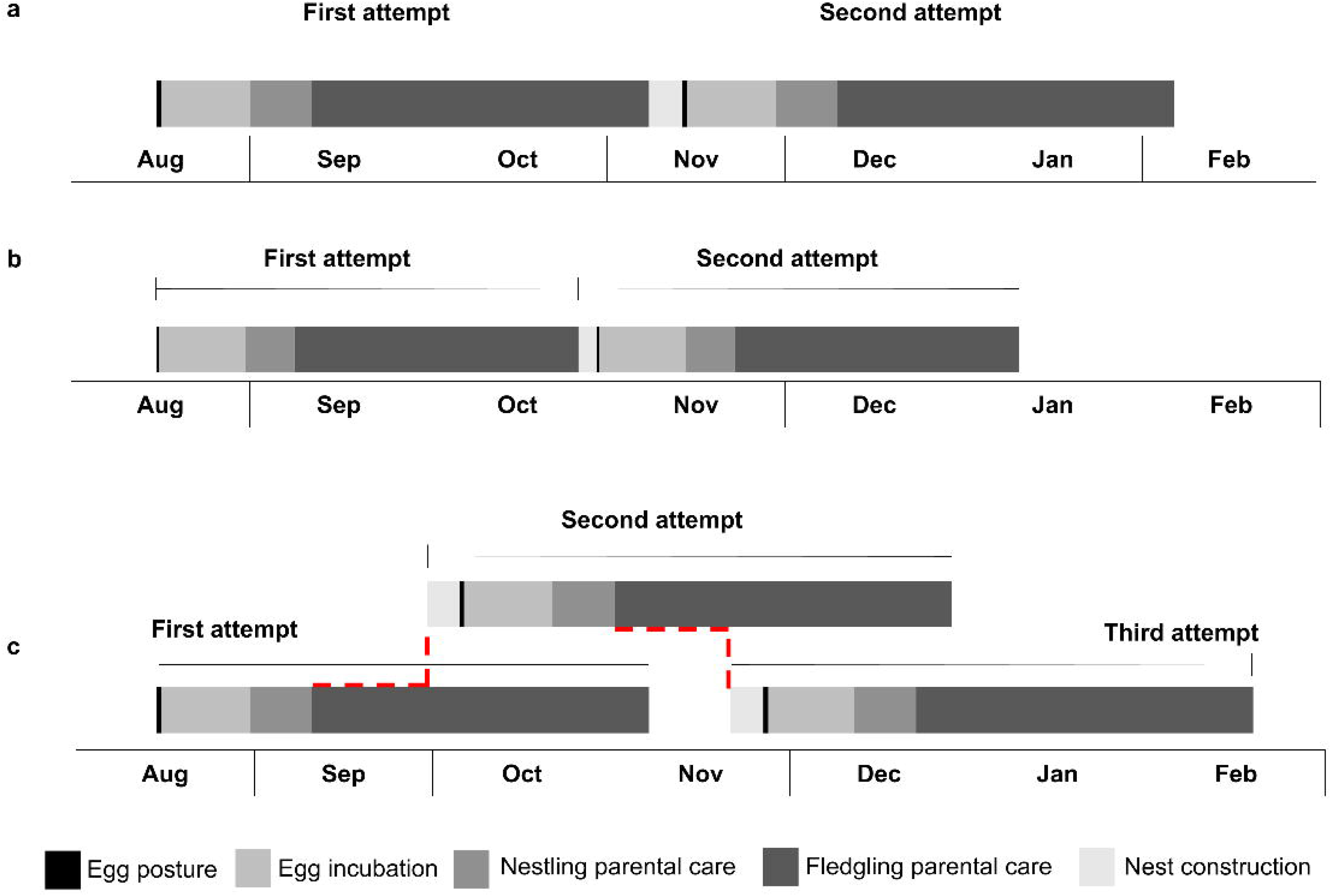
Hypothetical reproductive calendar of *Formicivora acutirostris* considering all the reproductive phenology of the species (Figure 4) and broods successfully producing fledgling that reach independence. In (a), we consider the average duration and in (b) the minimum duration for each of the stages of reproduction. In (c) we hypothesized three broods with production of at least one fledgling that reach independence, considering the average duration of each of the nesting and overlapping stages of broods from the 13^th^ day of life of the fledgling (s) (indicated in dashed red). Although we recorded egg laying in early February, they only occurred when there was no brood with production of at least one fledgling that reached independence in the previous months. The phenology starts from the date of the first egg laid ever recorded (August 14). The average and minimum duration of each of the stages of reproduction, in days, are: nest construction =7.95 / 4.0; laying = 1.69 / 1.0; incubation of eggs = 16.0 / 14.0; parental care of the nestlings = 10.0 / 8.0; and parental care of the fledglings = 56.7 / 48.0

## Discussion

### Functions associated with brood division in Formicivora acutirostris

Although nestlings and fledglings of *F. acutirostris* presented no appreciable differences in size and plumage, there was a non-random selection of the fledgling sex by the caring adult, suggesting a sex-biased selection of the fledgling (*pro* hypothesis #2). The 1) predation risk; 2) spatial segregation of siblings, 3) defensive behaviors of parents, and 4) low mortality of fledglings suggest that the permanent division of parental care from the first day of the fledgling phase is a behavioral strategy that reduces predation (*pro* hypothesis #1). In addition, the long-term brood division, fledgling recognition of the call of their adult, and improved the efficiency of the adult by caring for only one fledgling, render support to the social specialization and improvement in parental care efficiency as additional functions of brood division in the species (*pro* hypotheses #6 and #5, respectively).

Due to the fact that there are defensive behaviors from the parents aimed at fledglings at risk of drowning, together with the fact that drowning is a cause of mortality as soon as they leave the nests, we propose the division of parental care as a behavioral strategy that could increase the survival of the offspring reducing their drowning risk (drowning reduction hypothesis #7). Because the adults take the siblings to different places of the territory, which has resources partially and temporarily unavailable during the daily high tides, we also propose that the brood division in *F. acutirostris* is a strategy that reduces the food dispute among family members in its floodable environment (spatial division of food resource hypothesis #8). Finally, we suggest that the brood division is a strategy that allows concomitant reproduction thus increasing offspring productivity (concomitant reproduction hypothesis #9).

Since brood care is equivalent between parents and fledglings leave the nest together, the selection of parents by the siblings is not applicable to the brood division in *F. acutirostris* (*contra* hypothesis #3). Since we did not observe aggression between nestlings before the division, the competition between siblings seems not to apply to the species (*contra* hypothesis #4).

### Brood division in a marsh-dwelling bird

We verified brood division in all monitored *F. acutirostris* successful nests. The division involves sex selection, with males taking care of males and females taking care of females. This sex-matching division can be explained by the cultural transmission hypothesis, which states that foraging behaviors are learned and passed between parents and offspring of the same sex (McLaughin & Montgomeri 1985). In the case of *F. acutirostris*, the male is larger and has a larger beak, characteristics that make the foraging behavior between males and females slightly different (Reinert 2008), and that could justify the sex selection of the parents (hypothesis #2). We do not know how the recognition of the offspring sex occurs, but it may be due to subtle characteristics of sexual dimorphism that we cannot perceive (Vega et al. 2007). This raises doubt about the brood division in the congeneric *F. erythronotos* being randomized by the adult who witnessed a particular fledgling leaving the nest (Mendonça 2001). The possibility that there was a previous selection for the sex of the fledgling that the adult followed is not excluded.

*Formicivora acutirostris* presents a long-term brood division, but pressures of predation and drowning on its fledglings occur mainly in the first days after leaving the nest, contradicting the assumption that division could be limited only to the period of greatest pressure of loss of fledglings (Moreno 1984). As in *F. erythronotos*, the brood division in *F. acutirostris* occurs after the fledgling leaves the nest, unlike *T. atrinucha* (Mendonça 2001; Tarwater and Brawn 2008).

The brood division in *F. acutirostris* is a strategy that reduces offspring predation, which is the biggest cause of reproductive failure in tropical birds (Oniki 1979; Winkler 2016), and drowning. Contributing to reducing predation, the brood division improves the efficiency of parental care (hypothesis # 5), which is one of the main hypotheses for the brood division in birds (Smith 1978; Moreno 1984; McLaughlin and Montgomerie 1985; Anthonisen et al. 1997; Tarwater and Brown 2008). The exclusive care of a fledgling decreases the parent’s expenditure of energy and time to feed the fledgling, increasing the efficiency of care (Smith 1978). Similarly, social specialization contributes to these purposes (hypothesis #6), rendering the possibility of mutual learning of individual characters between parents and fledglings (Leedman and Magrath 2003). This would facilitate sound communication widely used by *F. acutirostris* to find the fledgling and protect him from danger.

The spatial division of food resources (hypothesis #8), a temporal division of the food niche, would reduce competition for food in an environment temporarily reduced by flooding. Usually, half or three-quarters of the vegetation is submerged at high tides. In addition, the habitat of the species is made of fragile pioneer formations that easily fall by the wind, which can affect vast areas of vegetation (Reinert et al. 2007), being another indication of the benefit that the spatial division of food resources would bring to *F. acutirostris*. This hypothesis can be confirmed by future works using kernel maps of the foraging area of the parents and their fledgling compared to the non-reproductive period.

The last of the additional hypotheses (#9) proposed to explain the brood division of *F. acutirostris* is related to increased parent productivity, with concomitant reproduction. Concomitant reproduction occurring in a breeding season that covers the entire reproductive phenology of the species (Figure 5) can lead to a third brood with fledglings reaching independence. This was observed only once in *F. acutirostris*, but it can raise the productivity of a breeding pair of up to six individuals in a single reproductive season (see Bornschein et al. 2015). Due to the low number of concomitant reproductions observed, it seems reasonable to assume that they occur in conditions of good nutrition of the parents and young and, furthermore, that they are favored when the current brood is of only one fledgling. Reproductive behaviors performed by the free parent may influence the other to split the parental care time with a new reproduction. Additionally, as demonstrated, there is an improvement in the efficiency of parental care (hypothesis #5) and greater efficiency in contact between the parent and their fledglings (hypothesis #6) by the division of young between parents. These behaviors can increase the time of the parent to seek food and, consequently, nourish themselves better for an upcoming or concomitant reproduction.

**Figure 5.**
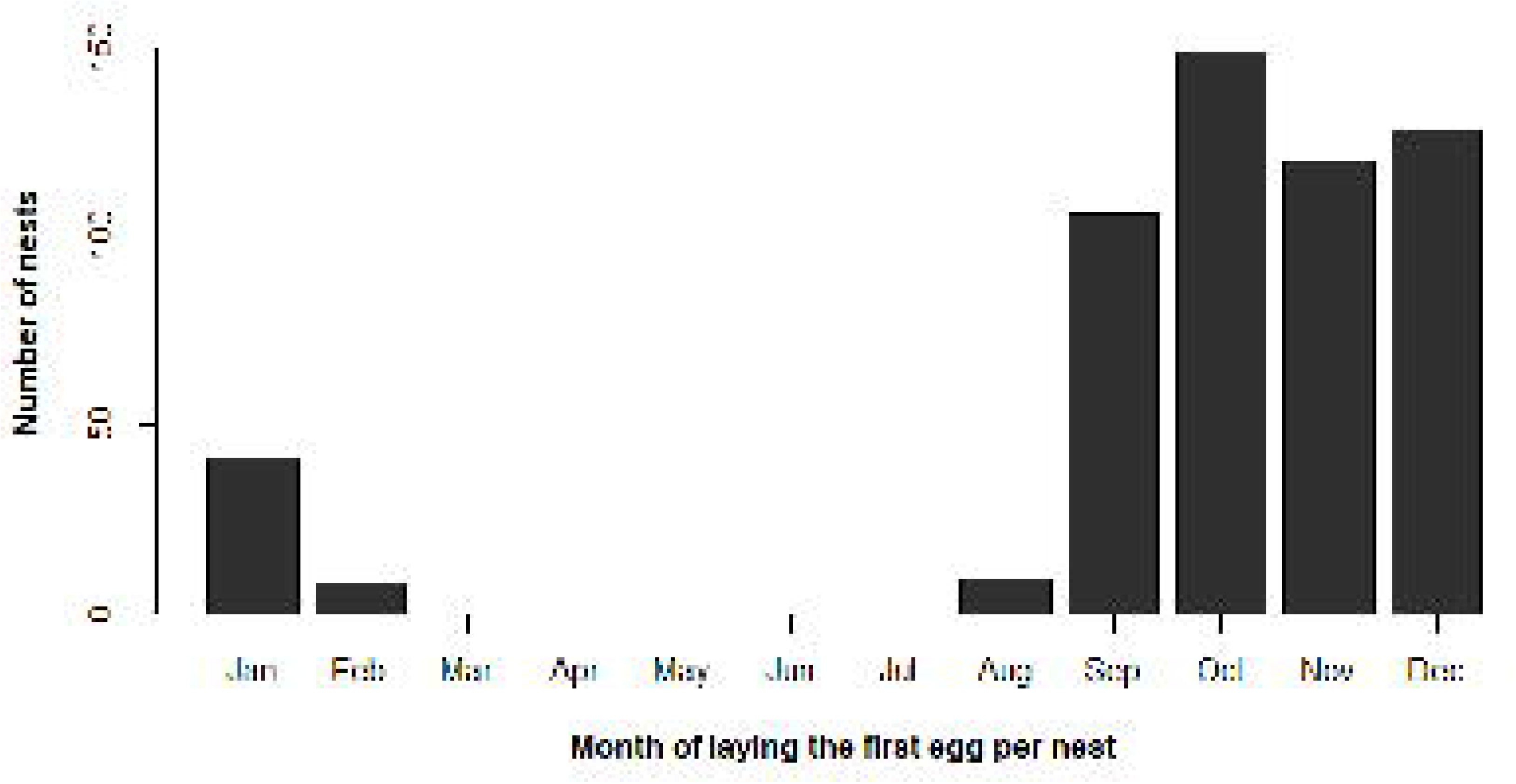
Reproductive phenology of *Formicivora acutirostris* in Guaratuba Bay, Paraná, southern Brazil (data from 2006 – 2022)

Some species optimize their reproductive success by increasing the number of young per reproduction, while others superimpose broods as a strategy to increase the number of breeding attempts (Burley 1980). Concomitant reproduction tends to occur mainly when the breeding season is short, the available food resource is abundant, and fewer young were generated in the previous attempt (Cate and Hilbers 1991; Blomqvist 2001; Hetmańsk and Wolk 2005). Concomitant reproduction still occurs mainly in species with long life expectancy and stable couples, which allow manipulation of parental care (Burley 1980). *Formicivora acutirostris* lives in permanent pairs and is a relatively long-lived species (Bornschein et al. 2015).

In conclusion, the associated functions of the brood division in *F. acutirostris* involve increased survival of the fledglings and the opportunity to generate more youngs. The brood division tends to increase reproductive success by increasing the chances of survival of the fledgling, as well as by increasing nesting attempts, provided by the improved care of a single fledgling. Considering that in addition to predation nests, nestlings and fledglings are also lost by flooding, we do not rule out the possibility that the concomitant reproduction in *F. acutirostris* also reflects a reproductive adaptation to “compensate” the reproductive losses by the limiting environment. The particular habitat of *F. acutirostris*, unusual for a Thamnophilidae, imposes restrictions for which there are behavioral responses by the species, as advocated by Tarwater and Brawn (2008). This raises the question of how well it can adapt to the perspective of severe and frequent floods that reduce productivity and temporarily limit the food resource in a climate change and sea level rise scenario. The continuity of the long-term study with *F. acutirostris* for the monitoring of brood division behavior and possibly consequent population dynamics is especially important for this unique Thamnophilidae.

## Acknowledgments

We thank the Fundação Grupo Boticário de Proteção à Natureza (FGBPN), Fundo Nacional do Meio Ambiente (FNMA), and Fundo Brasileiro para a Biodiversidade (FUNBIO), that financially supported long periods of research. State of São Paulo Research Foundation (FAPESP; process n° 2022/04847-7) and the National Council for Scientific and Technological Development (CNPq; process # 314038/2021-3) from the grant provided Giovanna Sandretti-Silva. Helena Zarantonieli, from Mater Natura - Instituto de Estudos Ambientais, that conducted the financial management of the projects. Leandro Corrêa, Carlos O. Gussoni, Tamiris Pereira Lima, and many others that assisted in the fieldwork. The analyzes were carried out as a result of the project “Olha o Clima, Litoral!” financed by the Petrobras Socioenvironmental Program

## Statements and Declarations

### Disclosure statemen

No potential conflict of interest was reported by the authors.

### Data availability

All data used for this study are available in the text and in the supplementary material.

