## Supplemental Table 1 for "Brood division in a marsh-dwelling bird and its relation with the increase in offspring survival, acceleration of renest, and reduced competition for food resources"

Table S**1.** Number of territories of *Formicivora acutirostris* monitored for the evaluation of parental care by reproductive season, Guaratuba Bay, Paraná, southern Brazil. Blank cells indicate the absence of monitoring in the respective reproductive season.

| Area | 2006 / 2007 | 2007 / 2008 | 2008 / 2009 | 2009 / 2010 | 2010 / 2011 | 2011 / 2012 | 2012 / 2013 | 2013 / 2014 | 2014 / 2015 | 2015 / 2016 | 2016 / 2017 | 2017 / 2018 | 2018 / 2019 | 2019 / 2020 | 2020 / 2021 | 2021 / 2022 |
| --- | --- | --- | --- | --- | --- | --- | --- | --- | --- | --- | --- | --- | --- | --- | --- | --- |
| Jundiaquara Island | 14 | 14 | 12 | 12 | 12 | 12 | 13 | 15 | 15 | 16 | 16 | 16 | 16 | 16 | 15 | 13 |
| Claro River |  | 5^a^ |  | 10 | 10 |  | 10 | 10 | 10 | 13 | 13 |  | 12 | 13 | 10 | 9 |
| Folharada Island |  |  |  | 10 | 10 | 13 | 13 | 12 | 13 | 11 | 12 |  | 11 | 11 | 9 | 8 |
| Lagoa do Parado |  |  |  |  |  |  | 5 |  |  |  |  |  |  |  |  |  |
| Total | 14 | 19 | 12 | 32 | 32 | 25 | 41 | 37 | 38 | 40 | 41 | 16 | 39 | 40 | 34 | 30 |

^a^ The area sampled was smaller at the time. It increased and remained standardized from the reproductive season of 2009/2010
