## Supplemental Table 2 for "Brood division in a marsh-dwelling bird and its relation with the increase in offspring survival, acceleration of renest, and reduced competition for food resources"

Table S**2.** Number of monitored nests and juveniles that achieved the independence of *Formicivora acutirostris* monitored for the evaluation of parental care by reproductive season, Guaratuba Bay, Paraná, southern Brazil. Blank cells indicate absence of monitoring in the respective reproductive season.

| Area | 2006 / 2007 | 2007 / 2008 | 2008 / 2009 | 2009 / 2010 | 2010 / 2011 | 2011 / 2012 | 2012 / 2013 | 2013 / 2014 | 2014 / 2015 | 2015 / 2016 | 2016 / 2017 | 2017 / 2018 | 2018 / 2019 | 2019 / 2020 | 2020 / 2021 | 2021 / 2022 |
| --- | --- | --- | --- | --- | --- | --- | --- | --- | --- | --- | --- | --- | --- | --- | --- | --- |
| Jundiaquara Island | 40 / 2 | 70 / 8 | 24 / 0 | 39 / 10 | 10 / 11 | 11 / 10 | 13 / 15 | 21 / 14 | 26 / 15 | 33 / 14 | 23 / 12 | 14 / 7 | 33 / 12 | 22 / 21 | 29 / 0 | 34 / 8 |
| Claro River |  | 9 / 0 |  | 30 / 0 | 25 / 2 |  | 15 / 4 | 16 / 10 | 17 / 5 | 17 / 9 | 10 / 4 |  | 4 / 3 | 16 / 10 | 6 / 0 | 19 / 6 |
| Folharada Island |  |  |  | 30 / 4 | 22 / 9 | 22 / 13 | 18 / 9 | 31 / 3 | 36 / 9 | 30 / 1 | 28 / 3 |  | 16 / 0 | 18 / 6 | 6 / 1 | 12 / 1 |
| Lagoa do Parado |  |  |  |  |  |  | 4 / 2 |  |  |  |  |  |  |  |  |  |
| Total | 40 / 2 | 79 / 8 | 24 / 0 | 99 / 14 | 57 / 22 | 33 / 23 | 50 / 30 | 68 / 27 | 79 / 29 | 80 / 24 | 61 / 19 | 14 / 7 | 53 / 15 | 56 / 37 | 41 / 1 | 65 / 15 |
